# A rotifer-derived paralytic compound prevents transmission of schistosomiasis to a mammalian host

**DOI:** 10.1101/426999

**Authors:** Jiarong Gao, Ning Yang, Fred A. Lewis, Peter Yau, James J. Collins, Jonathan V. Sweedler, Phillip A. Newmark

## Abstract

Schistosomes are parasitic flatworms that infect over 200 million people, causing the neglected tropical disease, schistosomiasis. A single drug, praziquantel, is used to treat schistosome infection. Limitations in mass drug administration programs and the emergence of schistosomiasis in non-tropical areas indicate the need for new strategies to prevent infection. It has been known for several decades that rotifers colonizing the schistosome’s snail intermediate host produce a water-soluble factor that paralyzes cercariae, the life-cycle stage infecting humans. In spite of its potential for preventing infection, the nature of this factor has remained obscure. Here, we report the purification and chemical characterization of Schistosome Paralysis Factor (SPF), a novel tetracyclic alkaloid produced by the rotifer *Rotaria rotatoria*. We show that this compound paralyzes schistosome cercariae and prevents infection, and does so more effectively than analogous compounds. This molecule provides new directions for understanding cercariae motility and new strategies for preventing schistosome infection.

## Introduction

Schistosomiasis – caused by parasitic flatworms of the genus *Schistosoma –* is a major neglected tropical disease, affecting over 200 million people, with over 700 million people at risk of infection (1–3). Praziquantel is currently the only drug used for treating schistosomiasis. Concerns about the emergence of drug resistance (4, 5) as well as limitations observed in mass drug administration programs (6–9) highlight the need to devise new strategies for preventing infection by these parasites. This need is amplified by the recent identification of people infected with human/livestock hybrid schistosomes and the geographical expansion of schistosomiasis to temperate regions (10–12).

Schistosomes have a complex life cycle that alternates between an intermediate host (snail) and a definitive host (mammal) via two free-living, water-borne forms called miracidia and cercariae, respectively (13) (Fig. 1A). For decades, inconsistency in cercarial production by snails and infectivity of mammalian hosts has been observed in most schistosome laboratories (14). Intriguingly, Stirewalt and Lewis reported that rotifer colonization on shells of the snail intermediate host (*Biomphalaria glabrata*) significantly reduced cercariae output, motility, and infectivity (15). Furthermore, they observed that cercarial motility was affected not only by the presence of rotifers, but also by rotifer-conditioned water, indicating that rotifers released water-soluble molecules with paralytic activity. Almost 40 years have passed since this important finding, yet this factor’s identity has remained a mystery.

**Fig. 1.**
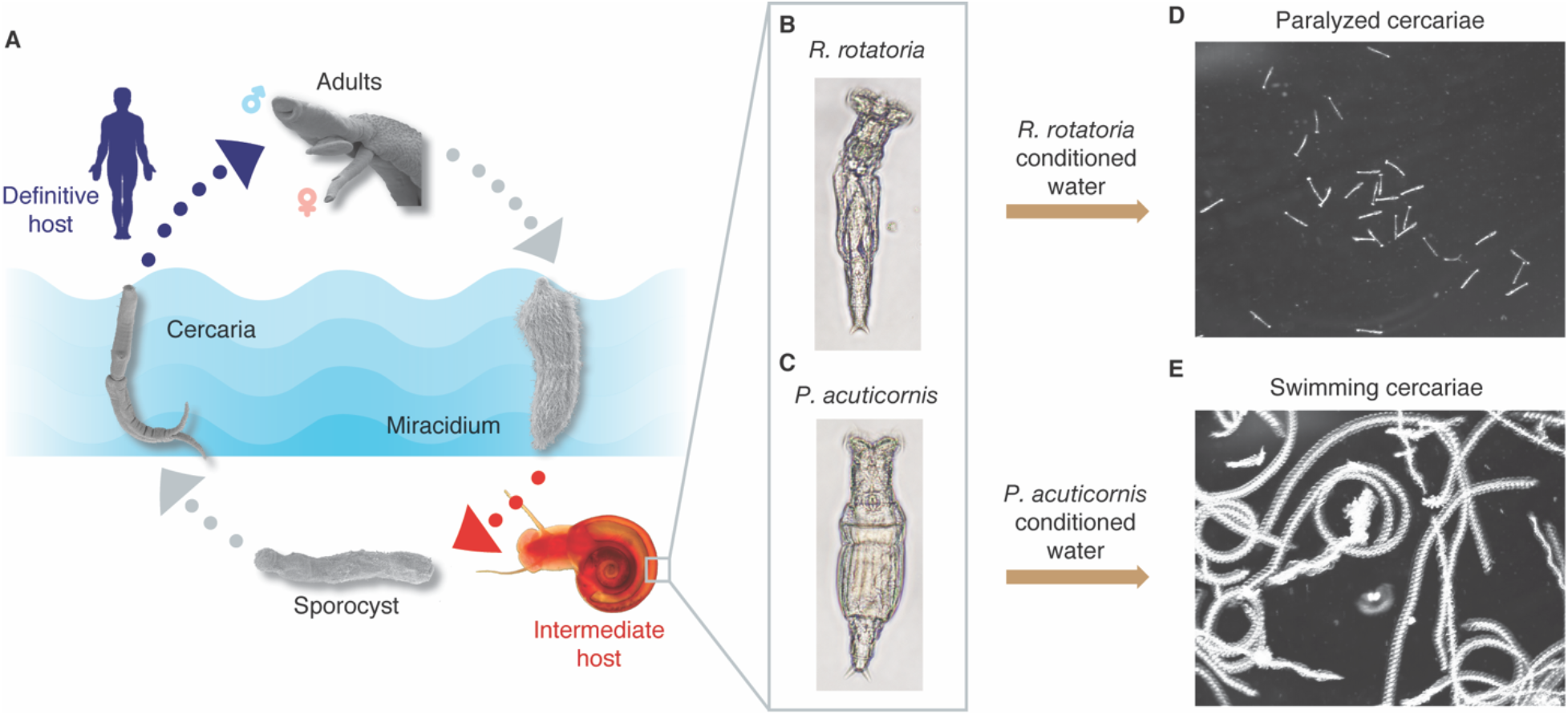
*R. rotatoria*-conditioned water paralyzes *S. mansoni* cercariae. (*A*) Life cycle of *S. mansoni*. Adult parasites, residing in the mammalian host vasculature, lay eggs (not shown). Upon exposure to fresh water, eggs release miracidia, which infect the appropriate snail host. Inside the snail the parasite reproduces asexually, ultimately producing large numbers of free-swimming infective larvae (cercariae) that can penetrate mammalian skin to continue the life cycle (adapted from (35)). (*B* and *C*), Bright-field microscopy images of *R. rotatoria* and *P. acuticornis*. (*D* and *E*) Maximum intensity projection (5 s, 150 frames) of cercariae motility after treatment with *R. rotatoria*- or *P. acuticornis*-conditioned water.

## Results and Discussion

### Purification of the rotifer-derived compound

Encouraged by this anti-cercarial effect and its potential to prevent schistosome infection, we sought to purify this paralyzing agent. We isolated individual rotifers from snail shells and found two species, *Rotaria rotatoria* (Fig. 1B) and *Philodina acuticornis* (Fig. 1C), as previously reported (15). To identify which rotifer was responsible for the paralytic effect, we grew clonal isolates of each species, producing rotifer-conditioned artificial pond water (APW). Adding *Rotaria*-conditioned APW to freshly collected cercariae resulted in gradual paralysis within five minutes (Fig. 1D). Most cercariae stopped swimming and sank to the bottom of the dish. Tapping the dish could stimulate their movement, but their response was limited to writhing on the dish bottom or short-distance swimming before becoming paralyzed again. In contrast, *P. acuticornis*-conditioned water had no effect (Fig. 1E).

To purify the paralyzing agent, we performed molecular weight cut-off filtration (MWCO) of rotifer-conditioned water and found that the activity was present in the <650 Da fraction. The <650 Da filtrate was fractionated by reversed-phase high-performance liquid chromatography (RP-HPLC) (Fig. 2A) and each fraction was tested qualitatively (i.e., plus/minus) for paralytic activity on cercariae. Paralysis was only observed following treatment with a peak eluting at 25-27 min (Fig. 2B). As expected, this peak was detected only in *R. rotatoria*-but not *P. acuticornis*-conditioned water (Fig. 2B). Performing a second round of HPLC on this active fraction and assaying all the resulting peaks revealed a single peak (eluting at 24-26 min) with paralytic activity (Fig. 2C). A predominant signal of m/z 273.16 (M+H) in this peak was revealed by matrix-assisted laser desorption/ionization mass spectrometry (MALDI-MS) (Fig. 2D). Consistent with the paralysis assay, this signal (m/z 273.16) was detected exclusively in the fraction eluting at 24-26 min but not in the fractions before or after (Fig. 2E). These results suggested that the component with m/z 273.16 was the paralyzing agent, which we named “Schistosome Paralysis Factor” (SPF). We then determined the monoisotopic mass for protonated SPF using high-resolution quadrupole time-of-flight (Q-TOF) MS, m/z 273.1595 (Fig. 2F), suggesting C_16_H_20_N_2_O_2_ as the best-fitting formula for SPF.

**Fig. 2.**
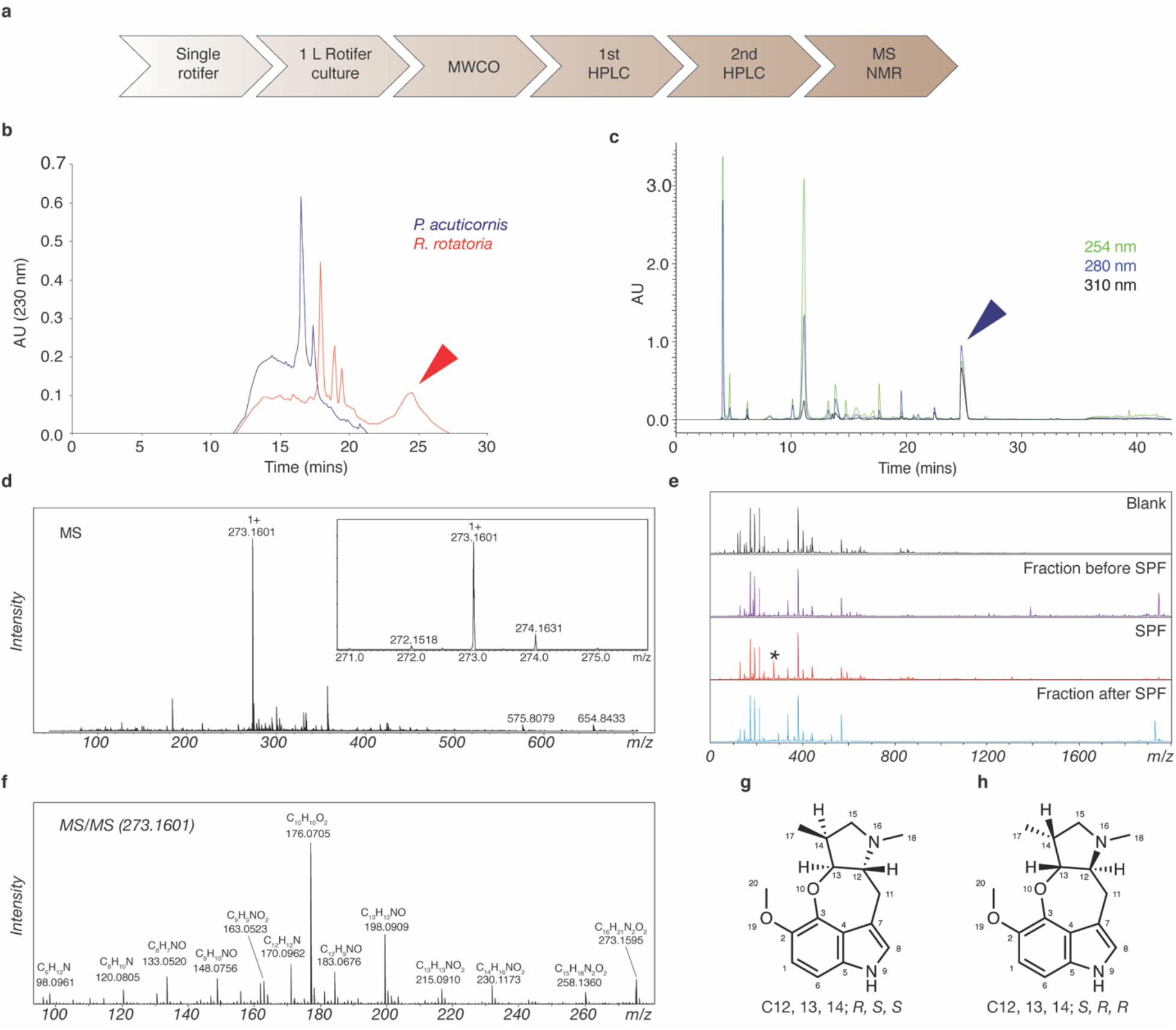
SPF is a novel tetracyclic alkaloid. (*A*) Flowchart for SPF purification. (*B*) 1^st^ HPLC plots of *R. rotatoria*- and *P. acuticornis*-conditioned water. All fractions were tested for bioactivity; the red arrowhead indicates the only active peak. (*C*) 2^nd^ HPLC plot of the bioactive fraction (red arrowhead in *B*). All peaks were tested for bio activity; the blue arrowhead indicates the only peak containing activity. (*D*) MS showing the dominant signal of m/z 273.1601 from the peak (blue arrowhead). (*E*) MS plots showing this signal (asterisk, m/z 273.1601) was only detected in the fraction eluting at 24-26 min. (*F*) Tandem MS acquired from high-resolution Q-TOF analysis. (*G* and *H*) NOESY resolved the relative stereochemistry of three chiral centers and narrowed it down to two possible configurations.

### SPF is a novel tetracyclic alkaloid

To elucidate its structure, we purified ~0.1 mg SPF from 25 L *R. rotatoria*-conditioned water. Nuclear magnetic resonance (NMR) spectroscopy revealed a novel tetracyclic structure. Briefly, ^1^H spectra showed the presence of 19 protons in the compound (Fig. S1), which agrees with the best-fitting formula and Hydrogen/Deuterium exchange mass spectrometry (MS) analysis (Fig. S2). Heteronuclear single quantum coherence spectroscopy (HSQC) revealed three methyl, two methylene, six methine groups, and five quaternary carbons (Fig. S3). Total correlation spectrometry (TOCSY) showed that aliphatic protons, except two methyl groups, are from one spin system (Fig. S4). The connectivity of the neighboring groups was derived from correlation spectroscopy (COSY) and heteronuclear multiple bond correlation (HMBC) spectra (Fig. S5 and S6). Overall, the aliphatic region is composed of a dimethylpyrrolidine structure which is linked to an indole via a CH_2_ group and an oxygen. Nuclear Overhauser effect spectroscopy (NOESY) suggested (*R, S, S*) or (*S, R, R*) configurations on the chiral centers (Fig. S7). Altogether, combined NMR analysis led to two possible structures (Fig. 2G and H; Table S1).

### SPF and its analogs paralyze cercariae in a dose-dependent manner

To test its dose dependency, we examined the paralytic effect of serially diluted SPF on cercariae by quantifying their movement over time. In the absence of SPF, over 82% of cercariae were free-swimming over three minutes (Fig. 3A). In 2.5 nM SPF, the percentage of free-swimming cercariae dropped to 67% at three-minutes post drug treatment. As the concentration of SPF increased, so did the rate of paralysis, and more cercariae were paralyzed at the end of treatment. We observed maximum effects in 250 nM and 2.5 μM SPF, with the majority of cercariae paralyzed within 30 s.

**Fig. 3.**
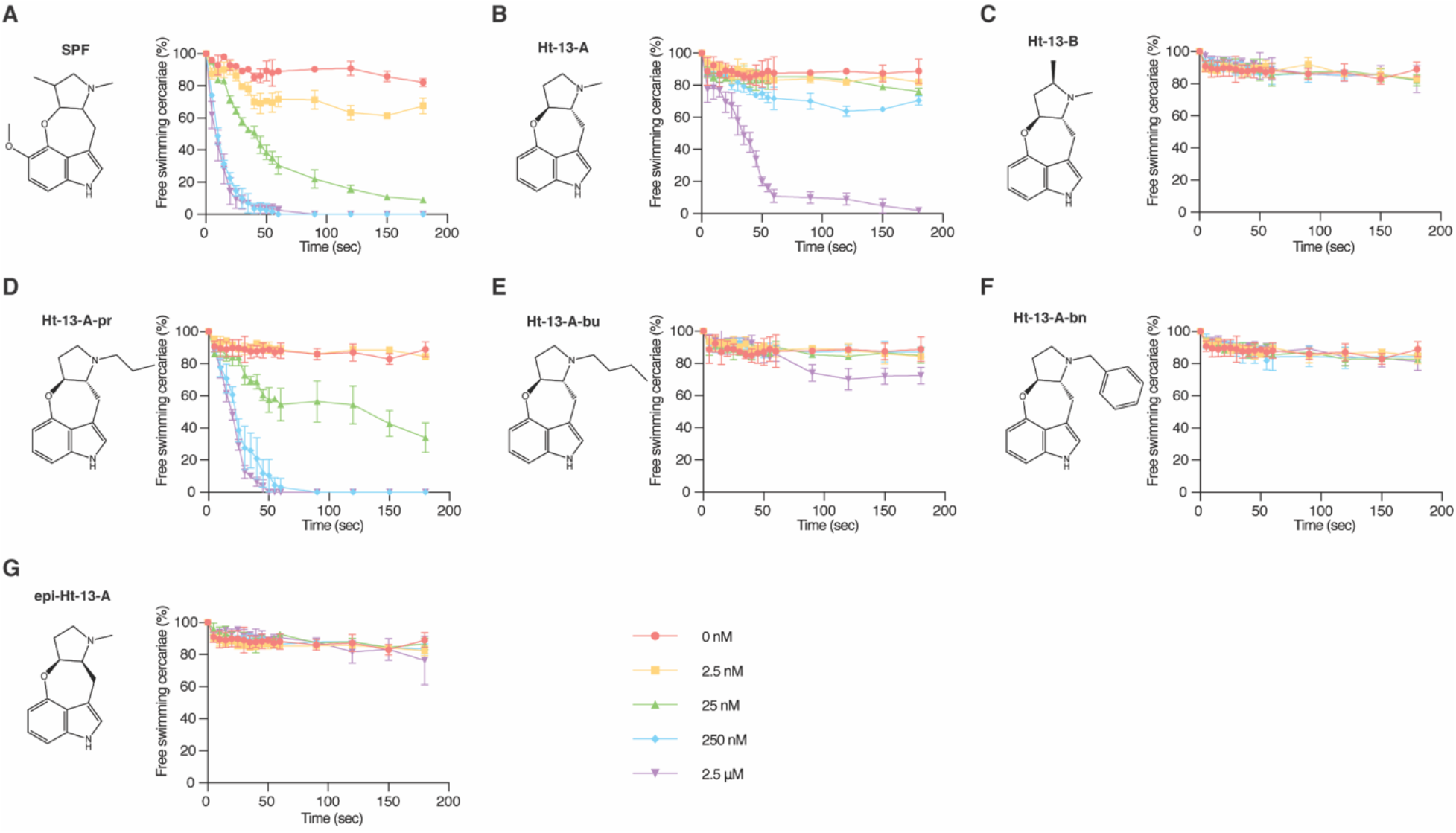
Structure-activity relationships of SPF and related compounds as measured by cercarial motility assays. (*A*-*G*), Percentage of cercariae (~50) continuing to swim over three minutes after addition of each compound at specified final concentrations. Triplicates were performed. Data are mean ± S.D.

Two natural compounds isolated from *Streptomyces sp.*, ht-13-A and ht-13-B (16), are structurally related to SPF. All three alkaloids share a novel oxepineindole framework fused with a pyrrolidine ring (Fig. 3A-C). Although synthesis of SPF has not been achieved, total syntheses of ht-13-A and ht-13-B have been reported (17–19). To test whether this shared tetracyclic scaffold is responsible for the paralytic effect, we analyzed structure-activity relationships by using ht-13-A, ht-13-B, three ht-13-A derivatives (18), and one epimer in cercarial paralysis assays. Importantly, ht-13-A, although not as potent as SPF, also had a paralytic effect on cercariae (Fig. 3B). In contrast, ht-13-B did not paralyze cercariae, suggesting that the extra methyl group disrupts interaction with the target (Fig. 3C). Of the three ht-13-A analogs, only ht-13-A-pr effectively paralyzed cercariae; it was more potent than ht-13-A, indicating that the nature of the side chain is important for proper target interaction (Fig. 3D-E). In contrast to ht-13-A, the epimer was unable to paralyze cercariae, supporting the (*R, S*) configuration of SPF at C12, 13 (Fig. 2G).

### SPF prevents mammalian infection

Since motility of the cercarial tail is essential for swimming and provides force for skin penetration (20–22), we examined whether SPF prevented infection. We treated ~200 cercariae with different concentrations of SPF for 10 mins, and tested their ability to infect mice after a 30-min exposure to their tails (N=6 for each condition). Six-weeks post infection, we euthanized the mice, counted schistosomes recovered after hepatic portal vein perfusion, and examined liver pathology. From controls, we recovered 83 adult worms on average (Fig. 4A), consistent with typical recoveries of ~40% (23). Livers from these mice appeared dark and contained extensive granulomas (Fig. 4A). In contrast, we did not recover any adult worms from mice after treatment with 250 nM or 2.5 μM SPF (Fig. 4B) and no granulomas were observed (Fig. 4A). Histological examination confirmed that these livers were free of schistosome eggs (Fig. 4E), suggesting complete inhibition of infection. These data are consistent with the full paralysis observed after treatment with 250 nM or 2.5 μM SPF (Fig. 3A). Although 25 nM SPF paralyzed most cercariae *in vitro*, the effects on mouse infection were not as severe (Fig. 4A). Mechanical and/or chemical stimuli from mouse tails may overcome SPF-induced paralytic effects at low SPF concentrations. Notably, neither Ht-13-A nor Ht-13-A-pr blocked infection as completely as 250 nM SPF, even at 25 μM (Fig. 4A, C, D, F, G). Under more realistic infection conditions, in which mouse tails were lifted 1-2 cm from the bottom of the test tube containing cercariae, so they had to swim actively towards the tail to infect the mouse, Ht-13-A and Ht-13-A-pr were still not as effective as SPF, which completely blocked infection (Fig. S8).

**Fig. 4.**
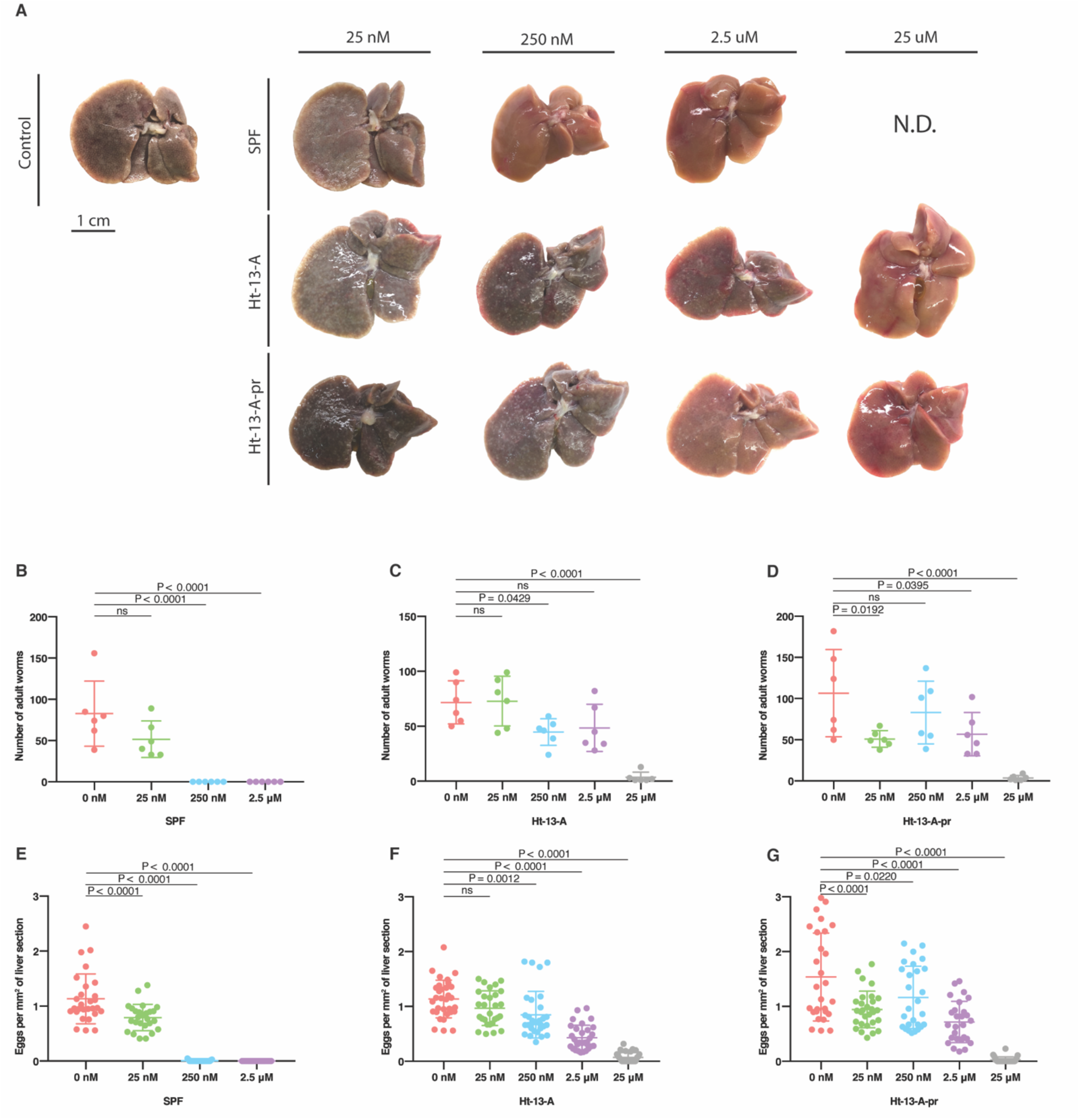
Treating cercariae with SPF, Ht-13-A or Ht-13-A-pr blocks schistosome infection and alleviates pathology. (*A*) Representative livers (post perfusion) from mice (N=6) exposed to drug-treated cercariae. Livers from mice treated with control and lower drug concentrations were darker in color and contained more granulomas (white spots). With higher drug concentrations, livers had normal morphologies with few or no granulomas. 25 μM SPF treatment was not determined (N.D.) due to limited amounts of purified SPF. (*B*-*D*) Numbers of adult worms recovered from exposed mice (two experiments for each drug, 6 mice total for each condition). (*E*-*G*) Numbers of schistosome eggs per area (/mm^2^) from liver sections (4-6 sections per mouse). Data (*B*-*G*) are mean ± S.D. Statistics: One-way ANOVA, post Dunnett’s test.

## Conclusion

This work has identified a novel tetracyclic alkaloid, produced by the rotifer *R. rotatoria*, that paralyzes the infective larvae of schistosomes. Although its mechanism of action remains unknown, its chemical structure provides important clues. SPF contains a serotonin backbone, suggesting that SPF might antagonize serotonin signaling, perhaps via G protein-coupled receptors (GPCRs) or serotonin-gated channels. Consistent with this idea, the structurally related compounds, ht-13-A and ht-13-B, bind several human serotonin receptors (16). In schistosomes, serotonin has been implicated in neuromuscular functions in multiple life stages (24–27); knocking down a serotonergic GPCR (Sm5HTR) in schistosomulae and adult worms led to decreased movement (28). Interestingly, praziquantel partially activates the human serotonin receptor, HT2BR, suggesting that it may also target schistosome serotonergic GPCRs (29).

The chemical ecology underlying *R. rotatoria*’s production of SPF is also unclear. Whether SPF is used naturally to combat other aquatic creatures (e.g., to prevent other rotifers from colonizing areas where *R. rotaria* live) and, thus, the effect on schistosome cercariae is indirect, or whether SPF benefits the rotifer’s commensal host will require further study. Given that the previously described related compounds are produced by *Streptomyces sp*.(16), it will also be important to examine the possibility that SPF is not directly produced the rotifer, but rather by constituent(s) of its own microbiome. In the past few decades, the discovery and development of natural products have helped combat parasitic diseases (30). Based on its ability to block infection, SPF holds great promise as an anti-schistosomal agent. Identifying the biologically active chemical scaffolds and understanding SPF’s mode of action are expected to provide important clues for preventing schistosomiasis.

## Materials and Methods

### Artificial pond water

Four stock solutions were prepared to make artificial pond water (31): 1) 0.25 g/L FeCl_3_ • 6H_2_O, 2) 12.9 g/L CaCl_2_ • 2H_2_O, 3) 10g/L MgSO_4_ • 7H_2_O, and 4) 34 g/L KH_2_PO_4_ 1.5 g/L (NH_4_)_2_SO_4_, pH 7.2. For 1L artificial pond water, we added 0.5 mL of FeCl_3_ solution, 2.5 mL CaCl_2_ solution, 2.5 mL MgSO_4_ solution and 1.25 mL phosphate buffer.

### Obtaining *S. mansoni* cercariae

Infected *B. glabrata* snails provided by Biomedical Research Institute (BRI, Rockville, MD) were maintained in artificial pond water and fed Layer Crumbles (chicken feed) (Rural King, Mattoon, IL). To obtain *S. mansoni* cercariae, *B. glabrata* snails were exposed to light at 26°C for 1-2 hrs. Artificial pond water containing cercariae was passed through 100 μm cell strainer (Falcon) to remove snail food and feces. Cercariae were then collected using custom-made 20 μm cell strainers.

### Rotifer culture

Since both rotifer species reproduce parthenogenetically, we clonally expanded each species into one-liter cultures from a single rotifer. Individual rotifers (*R. rotatoria* and *P. acuticornis*) were initially isolated from the shell of *B. glabrata* and cultured in artificial pond water in 24-well plates. Each individual colony was expanded into ever-larger culture volumes and ultimately maintained in two-liter flasks. Both species were fed Roti-rich liquid invertebrate food (Florida Aqua Farms Inc). Rotifer-conditioned water was collected every month by filtering out the rotifers using a 20 μm cell strainer. Conditioned water from *R. rotatoria* paralyzed cercariae completely within 3-5 minutes, resembling treatment with 25 nM purified SPF, and suggesting an SPF concentration in the range of tens of nM. Filtered rotifers were then passaged to fresh artificial pond water to propagate the cultures.

### Crude rotifer-conditioned water preparation

One liter rotifer media was lyophilized, reconstituted with 50 mL dH_2_O and filtered through 10,000 and 650 (MWCO) Pall Minimate TFF Capsules with Omega membrane (Ann Arbor, MI). Filtrate (<650 Da) was freeze dried. For RP-HPLC, 300 mg of the dried material was dissolved in dH_2_O and run on a RP-HPLC – Merck Chromolith semi-prep RP-18e column (Darmstadt, Germany) at 5 ml/min using a gradient of 100% A to 60% B in 60 min. 10 mL fractions were collected and assayed for biological activity. Fractions containing biological activity were saved for further study.

### Further purification of rotifer media

The bioactive fractions were pooled, freeze dried with SpeedVac (Savant, MA), reconstituted with 500 μL dH_2_O and injected into a 4.6mm diameter × 25cm Symmetry column (Waters, MA). Breeze2 analytical LC system (Waters, MA) was employed for separation at 0.5 ml/min with the following solvents and gradients: Solvent A, 0.1% formic acid (FA); solvent B, methanol with 0.1% FA; 0—10 min 0—10% B, 10—30 min 10—35% B, 30—33 min 35—80% B, 33—37 min 80—80% B, 37—40 min 80—0% B. Eluents were collected manually based on peak elution. All fractions were lyophilized, reconstituted with water and analyzed with MALDI-MS. Fractions containing biological activity were saved for future use.

### Matrix-assisted laser desorption/ionization mass spectrometry (MALDI-MS) analysis

For each collected fraction, 1 µL of sample solution was spotted on ground steel MALDI target and mixed with 1 µL of alpha-cyano-4-hydroxy-cinnamic acid (CHCA, Sigma-Aldrich, MO) solution (10 mg/mL CHCA in 50% acetonitrile solution with 0.005% trifluoroacetic acid). Mass calibration, spectra acquisition and analysis were performed under conditions as previously described (32).

### High-resolution quadrupole time-of-flight mass spectrometry (Q-TOF MS) analysis

1 µL of the bioactive fraction was separated on a Magic 0.1 × 150mm column (Michrom, CA) and analyzed with maXis 4G mass spectrometer (Bruker, MA) using previously established methods for metabolite study (33). The separation was performed at 300 nl/min by use of solvent A (95% water, 5% acetonitrile with 0.1% FA) and solvent B (5% water, 95% acetonitrile with 0.1% FA) with the following gradient conditions: 0—5 min 4% B, 5—50 min 4—50% B, 50—52 min 50—90% B, 52—60 min 90% B, 60—70 min 90—4% B, 70—90 min 4% B.

### Hydrogen/deuterium (H/D) exchange analysis

Acidified deuterated methanol (CD_3_OD, methanol—d4, Sigma—Aldrich, MO) was made by adding 1 µL of deuterated FA into 1 mL of CD_3_OD. 2 µL of the bioactive fractions were added into 18 µL of acidified methanol above. 15 µL of the mixture were analyzed by direct infusion into a modified 11 Tesla FTMS (Thermo Scientific, MA) through NanoMate robot (Advion, NY) (34). Full spectra were acquired with resolution set at 100k.

### Nuclear magnetic resonance (NMR) analysis

Purified bioactive materials were dissolved in 250 µL of CD_3_OD and transferred into a 5 mm Shigemi NMR tube with a glass magnetic plug with susceptibility matched to CD_3_OD on the bottom. All NMR data were collected at 40°C on an Agilent VNMRS 750 MHz spectrometer equipped with a 5 mm Varian indirect detection probe with z gradient capability. Collected NMR data included 1H spectrum, gradient selected correlation spectroscopy (gCOSY), total correlation spectroscopy (TOCSY), nuclear Overhauser enhancement spectroscopy (NOESY) with a mixing time of 500 ms, heteronuclear single quantum coherence spectroscopy (^1^H—^13^C HSQC) and heteronuclear multiple-bond correlation spectroscopy (^1^H—^13^C HMBC). The NMR spectra were analyzed using Mnova NMR software (Mestrelab Research, Spain).

### Determination of SPF concentration

The proton quantification experiments were performed at 23°C on an Agilent 750 MHz VNMRS NMR spectrometer equipped with a 5 mm triple-resonance (^1^H/^13^C/^15^N) indirect-detection probe with XYZ PFG gradient capability. The probe was calibrated using the qEstimate tool in the Agilent VnmrJ4.2 software with a known standard. The proton spectrum of the sample was collected with a 90-degree pulse angle of 8.5 ms, 16 scans and 10.4 s delay between scans. The Agilent VnmrJ4.2 software was used to determine the concentration of the sample based on the integration values of proton peaks. A total of 5 well-resolved proton peaks (7.12ppm (1H), ~6.89ppm (2H), 4.41 (1H), 3.83 (3H), and ~3.58 (2H)) was used, and the concentration of the sample was 1.55 ± 0.07 mM. All concentrations used in the cercarial paralysis assay were calculated based on this value.

### Cercarial paralysis assay

To capture the whole field while avoiding excess reflected light in a well, we used the lid of 96-well plate (Costar). 40 μL of artificial pond water containing ~50 cercariae were added to each shallow well on the lid. 10 μL of SPF (dissolved in APW) was then added to reach the final concentration indicated. Using a high-speed camera (Olympus i-SPEED TR), attached to a stereomicroscope (Leica MZ125), we recorded cercariae movement at 20-60 fps at 1.25X magnification just prior to addition of test compounds until 3-4 min after treatment started. Raw movies were converted to. avi files using i-SPEED Viewer and compressed into JPEG format using ImageJ (addition of compound is considered time 0). We then counted the numbers of free swimming/paralyzed cercariae every 5 s for 1 min and every 30 s thereafter for 3 min. The number of dead cercariae (those that never swim before and after SPF treatment) were subtracted from data. Experiments were performed in biological triplicate.

### Mouse infectivity assay

Mouse infections were performed by exposing mouse tails to *S. mansoni* cercariae according to standard protocol from BRI(23) with slight modifications. Briefly, we secured mice in rodent restrainers (Thomas Scientific, Cat #551-BSRR) and put them vertically on top of a rack with grids. We pipetted 100 μL of each drug at proper concentration into a skinny glass tube (Fisher Scientific, Cat #14-958A) inside a 12 × 75 mm holding glass tube (VWR, Cat # 47729-570). 300 μL of APW containing ~200 cercariae were pipetted into each skinny tube and incubated for 10 mins before we inserted the mouse tail. Mouse tails were wiped with APW-moistened Kimwipes, inserted into the skinny tube, and exposed to cercariae for 30 mins. The mouse tail was touching the bottom of the test tube unless otherwise specified. We euthanized and perfused these mice six week-post infection according to standard protocols(23). For each drug, we initially used three mice for controls (APW only) and three mice for each concentration tested except for 25 nM Ht-13-A and Ht-13-A-pr. We then repeated the experiments again with three mice for each condition. In addition to that, we included six mice for 25 nM Ht-13-A and Ht-13-A-pr.

Adult worms were recovered by hepatic portal vein perfusion and briefly incubated in 2.5% Tricaine (Sigma) to separate males and females. We counted total numbers of adult worms under a stereomicroscope (Leica MZ75). Livers from infected mice were fixed in 4% formaldehyde in PBS overnight. Largest liver lobes (left lobe) were submitted to University of Wisconsin-Madison Histology Core Facility for sectioning and Hematoxylin and Eosin staining. Each left lobe was evenly cut into 4-6 pieces and paraffin embedded on a large cassette. One slide (4-6 liver sections) for each liver was used for histological examination, which provided a representative view throughout the whole liver lobe. We took a tiled image of the whole slide using a Zeiss Axio Zoom microscope and used ImageJ to determine the area of each section. Total numbers of eggs in each section were counted and normalized to the area.

In adherence to the Animal Welfare Act and the Public Health Service Policy on Humane Care and Use of Laboratory Animals, all experiments with and care of mice were performed in accordance with protocols approved by the Institutional Animal Care and Use Committee (IACUC) of the University of Wisconsin-Madison (protocol approval number M005569).

### Statistical analysis

GraphPad Prism (Version 7) was used for all statistical analyses. One-way ANOVA test followed by Dunnett’s multiple comparison test was used. Mean ± S.D. is shown in all figures.

## Acknowledgments

*B. glabrata* snails were provided by the NIAID Schistosomiasis Resource Center of the Biomedical Research Institute (Rockville, MD) through NIH-NIAID Contract HHSN272201700014I for distribution through BEI Resources. We thank: Melanie Issigonis, Umair Khan, Jayhun Lee, and Tania Rozario for helpful discussions and comments on the manuscript; Tracy Chong and Jayhun Lee for help maintaining the schistosome life cycle; Melanie Issigonis for solving the pond water crisis; Björn Söderberg and Yanxing Jia for providing Ht-13-A, -B, and derivatives; Lingyang Zhu for expert assistance with NMR; Brian Imai for assistance with SPF purification; as well as Peg Stirewalt, James Leef, Tom Nerad, and Paul Mazzocchi for their early efforts to help solve this puzzle.

