## Supplemental data for "A rotifer-derived paralytic compound prevents transmission of schistosomiasis to a mammalian host"

**Supporting information**

Table of Contents……………………………………………………………………...…………22

Supplementary Figures S1-8

Supplementary Figure S1.
$^{1}$H Nuclear magnetic resonance (NMR) spectrum of SPF……………………………….23

Supplementary Figure S2.
Fourier-transform mass spectrometry (FTMS) of SPF before and after deuterium exchange……………………………………………………………………………………24

Supplementary Figure S3.
$^{1}$H—$^{13}$C Heteronuclear single quantum coherence spectroscopy (HSQC) NMR spectrum of
SPF…………………………………………………………………………………......25

Supplementary Figure S4.
Total correlation spectrometry (TOCSY) NMR spectrum of SPF……………………....26

Supplementary Figure S5.
Correlation spectroscopy (COSY) NMR spectrum of SPF……………………………...27

Supplementary Figure S6.
H—$^{13}$C Heteronuclear multiple bond correlation (HMBC) NMR spectrum of SPF….....28

Supplementary Figure S7.
Nuclear Overhauser effect spectroscopy (NOESY) NMR spectrum of SPF………….....30

Supplementary Figure S8.
Stringent mouse infection experiment…………………………………………………...32

Supplementary Table S1.
Summary of protons and carbons from $^{1}$H, COSY, HSQC, HMBC and NOESY………33

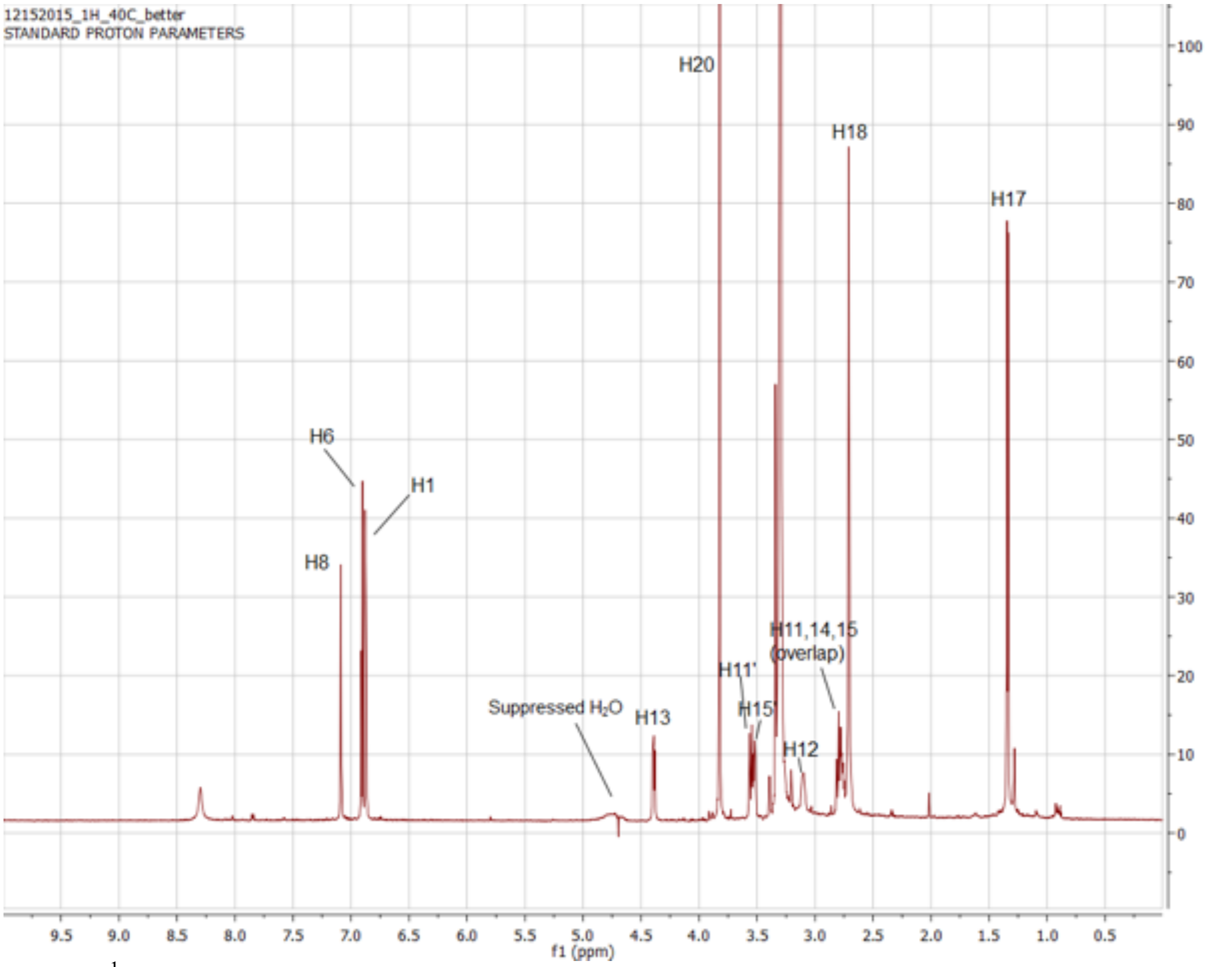


Fig. S1. [1]H Nuclear magnetic resonance (NMR) spectrum of SPF. Peak areas of the non-overlapping peaks were integrated and protons (δH 1.34, 3.10, 3.52, 3.56, 3.81, 4.40, 6.86, 6.90 and 7.09) showed integer ratios, supporting the mass spectrometry results that their signals were from the same compound. After adding the integration of overlapping peaks (δH 2.70, 2.72, 2.77, 2.79), a total of 19 protons were discovered, consistent with the best-fitting formula from the mass spectrometry results: ($C_{16}H_{20}N_2O_2$).

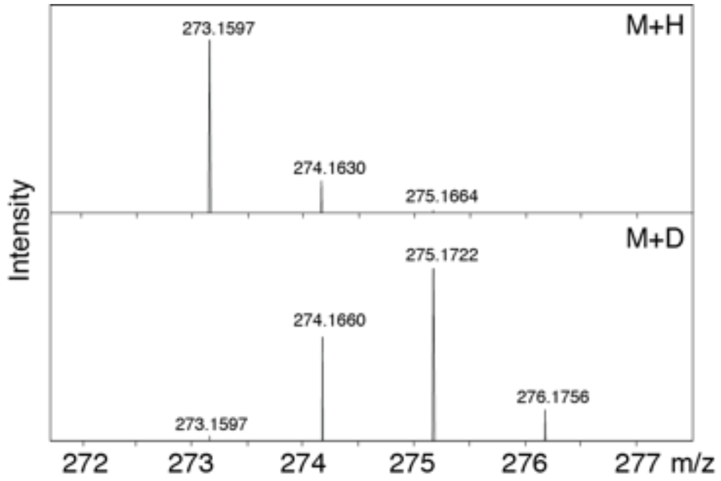

Fig. S2. Fourier-transform mass spectrometry (FTMS) determined the accurate *m/z* of the target molecule and revealed its isotopic pattern. Before deuterium exchange (top panel), 273.1597 was the measured *m/z* of the target molecule. After deuterium exchange (bottom panel), m/z of the base peak increased to 275.1722 (deuterium singly charged target molecule with one proton replaced by deuterium), suggesting the presence of one exchangeable proton in SPF.

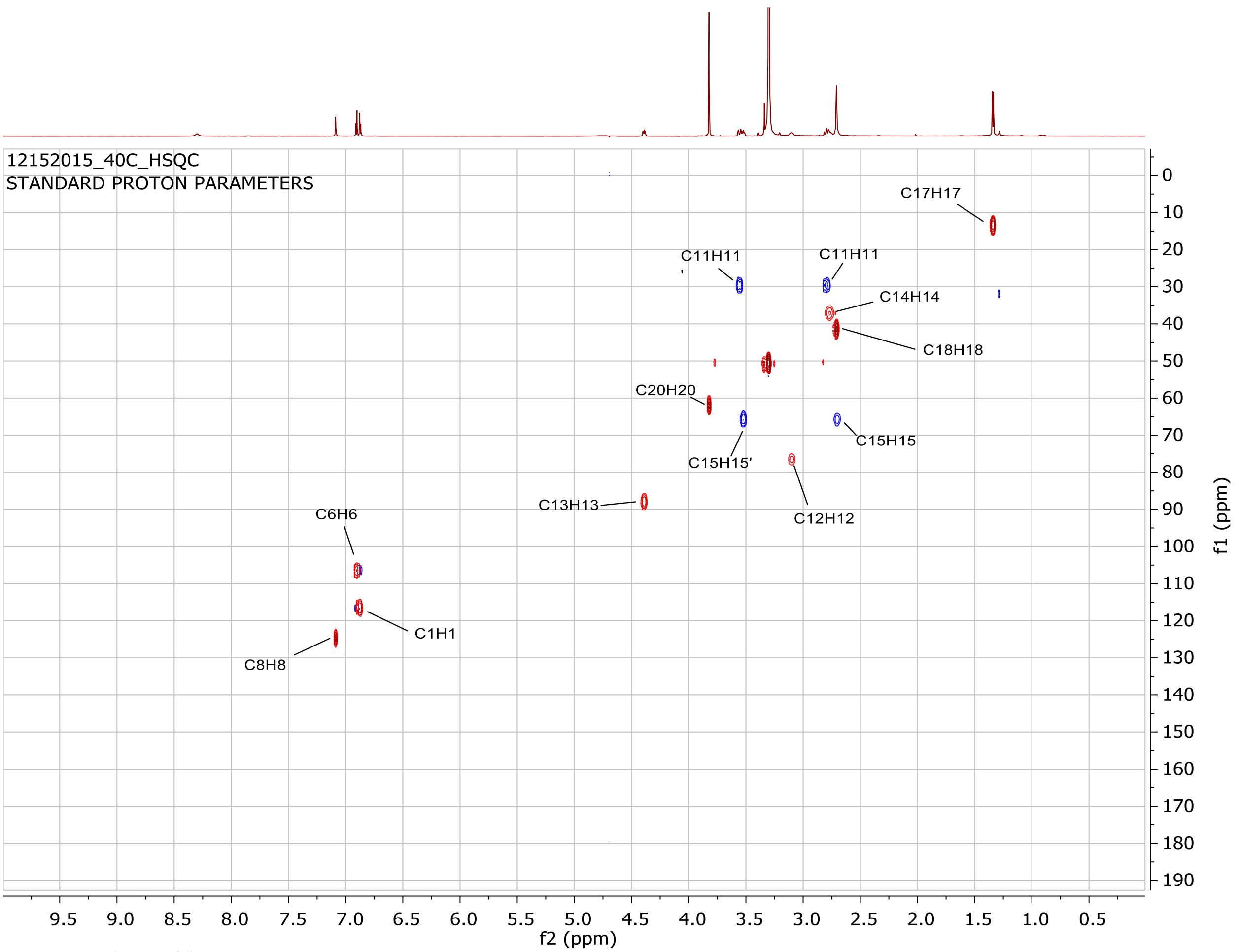


Fig. S3. $^{1}H$—$^{13}C$ Heteronuclear single quantum coherence spectroscopy (HSQC) NMR spectrum of SPF. HSQC revealed the cross—correlation between directly bonded proton and carbon nuclei and determined the number of methyl, methylene and methine groups. 19 protons were attached to 11 carbons, including: three methyl groups (δC 14.0, δH 1.34; δC 41.7, δH 2.71; δC 62.2, δH 3.81); two methylene groups (δC 29.0, δH 2.79, 3.56; δC 65.8, δH 2.70, 3.52); and six methine groups (δC 116.8, δH 6.86; δC 106.6, δH 6.90; δC 124.9, δH 7.09; δC 76.7, δH 3.10; δC 88.2, δH 4.40; δC 37.2, δH 2.77). The other 5 carbons that did not show up in the HSQC spectrum are the quaternary carbons. Based on the carbon chemical shift, the two methyl groups (δC 41.7, δH 2.71 and δC 62.2, δH 3.81) are likely to be bound to nitrogen and oxygen, respectively.

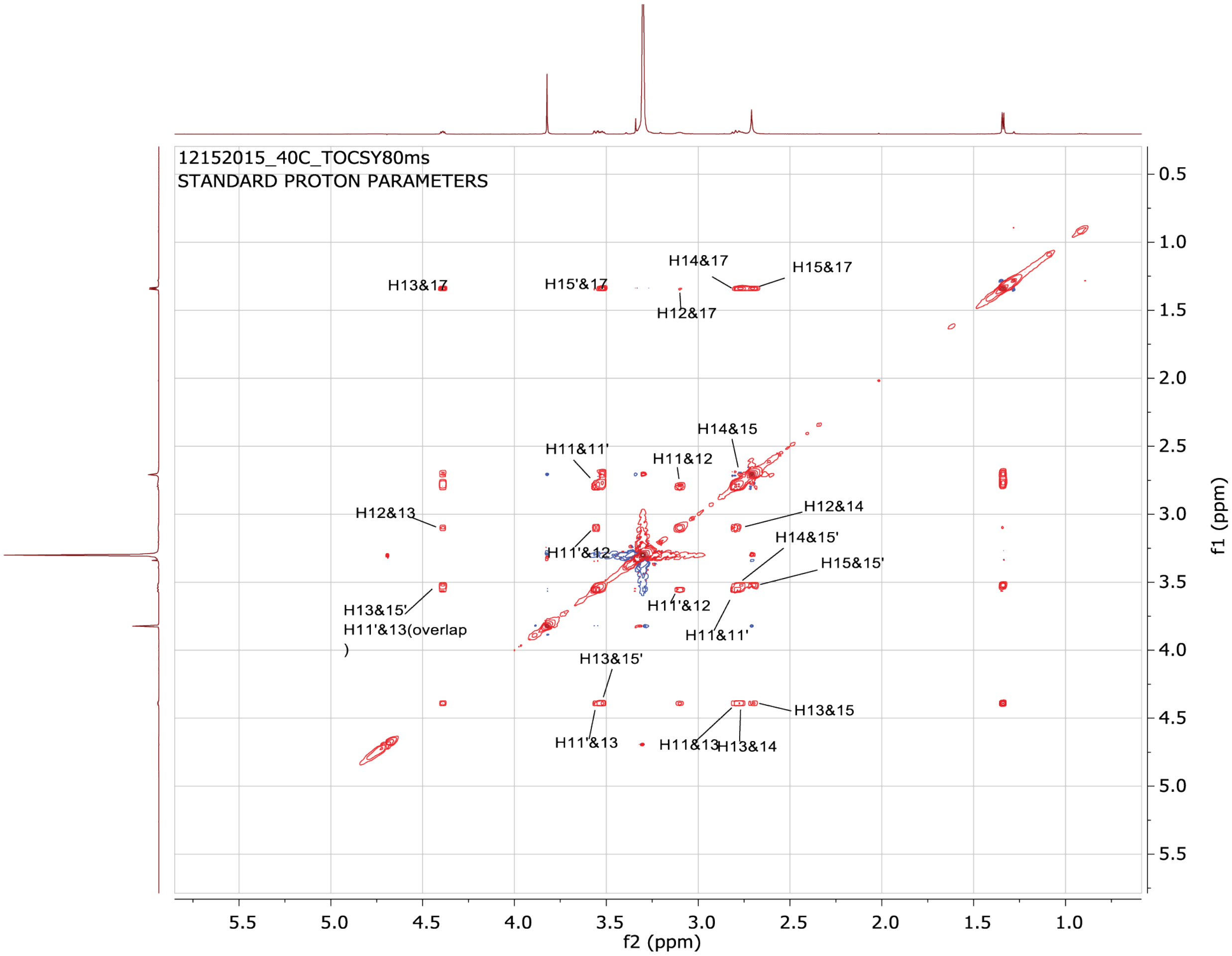


Fig. S4. Total correlation spectrometry (TOCSY) NMR spectrum of SPF. TOCSY revealed that the aliphatic protons except the two methyl groups (δH 2.71 and 3.81) found binding to N and O in HSQC (Fig. S3) are from a single spin system. Cross-peaks were also observed among the aromatic proton δH 7.09 and the aliphatic protons (δH 3.56, 2.79 and 3.10), due to long-range couplings.

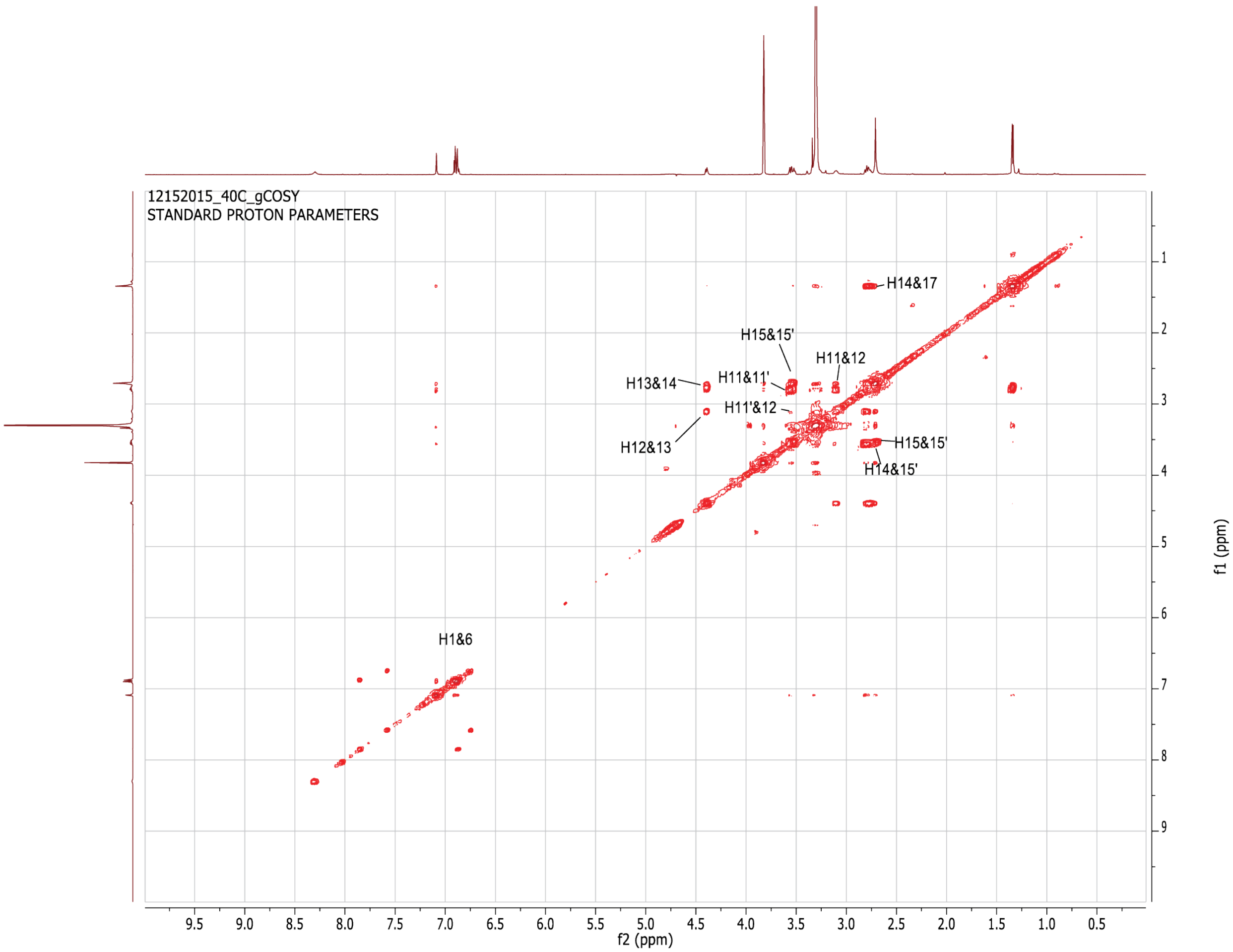


Fig. S5. Correlation spectroscopy (COSY) NMR spectrum of SPF. Based on HSQC, protons 11 and 11' (δH 2.79 and 3.56) are on the same carbon. Both have cross-peaks with proton 12 (δH 3.10) on COSY, which has an additional cross-peak with proton 13 (δH 4.40). This suggests $CH_2$ (C11, H11 and 11')-CH (C12, H12)-CH (C13, H13) connectivity. Similarly, proton 14 (δH 2.77) is connected to CH (C13, H13). Methyl group $CH_3$ (proton 17, δH 1.34) and $CH_2$ group (proton 15, 15', δH 2.70, 3.52) are directly connected to CH (proton 14).

A

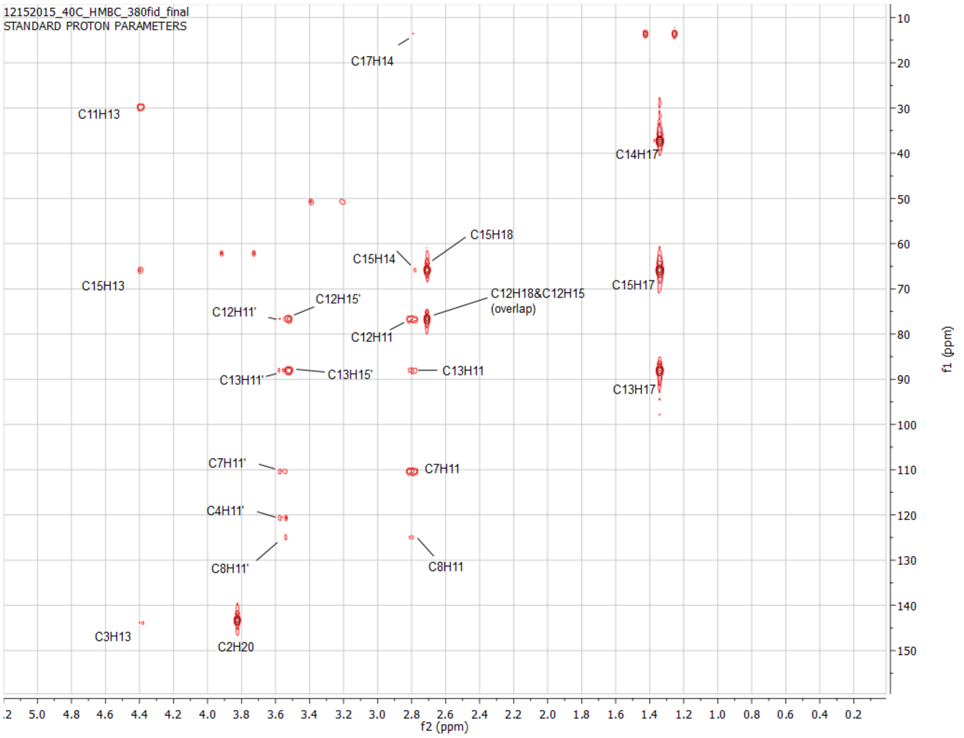


Fig. S6. H—[13]C Heteronuclear multiple bond correlation (HMBC) NMR spectrum of SPF. **(*A*) Aliphatic region.** Given the results from the COSY (Fig. S5) and the chemical shifts of C12 and C15 (δC 76.7 and 65.8), C12 and C15 are joined to a heteroatom. Since proton 18 (δH 2.71) has cross-peaks with both C12 and C15, it is a nitrogen atom that connects methyl group (δC 41.7, δH 2.71 on position 18), CH group (δC 76.7, δH 3.10) and $CH_2$ group (δC 65.8, δH 2.70 and 3.52). C13 has a chemical shift of 88.2 ppm, suggesting its connection to an oxygen. With HMBC, TOCSY, HSQC and COSY, the connectivity of the aliphatic portions are resolved.

B

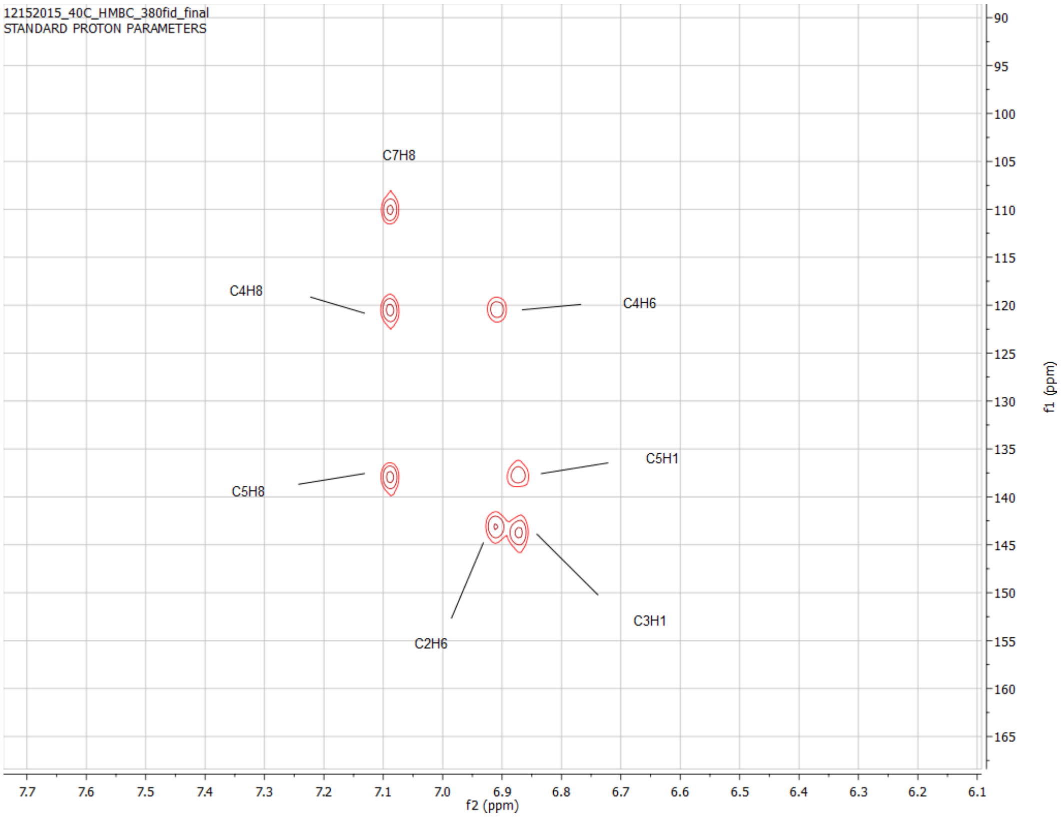


(***B***) **Aromatic region.** The connectivity-built aliphatic structure has the formula $C_7H_{13}NO$, which leaves $C_9H_6NO$ after subtracting from the best-fitting formula. HSQC (Fig. S3) showed the existence of methoxyl group (δC 62.2, δH 3.81). Therefore, the aromatic region was composed of $C_8H_3N$. HMBC data showed that three aromatic protons were located in different rings, implying a fused aromatic ring structure with one nitrogen. A substituted indole was the most common structure utilized in organisms with the matching formula. In addition, HMBC showed that protons on the methoxyl group (δH 3.81) and the aromatic proton (δH 6.90) have cross-peaks with carbon (δC 143.1), suggesting they are meta to each other. The other proton (δH 6.86) was vicinal to proton (δH 6.90) because of their coupling seen in the COSY spectrum (S5). The aromatic singlet proton δH 7.09 showed cross-peaks with three aromatic carbons, two of those carbons (δC 120.6 and δC 138.1) had cross-peaks with protons (δH 6.86 and δH 6.90) respectively, consistent with an indole configuration. HMBC further confirmed C (δC 110.6) was linked to $CH_2$ (δH 2.79 and 3.56), and C (δC 143.7) was linked to the CH (δC 88.2, δH 4.40) across an oxygen atom.

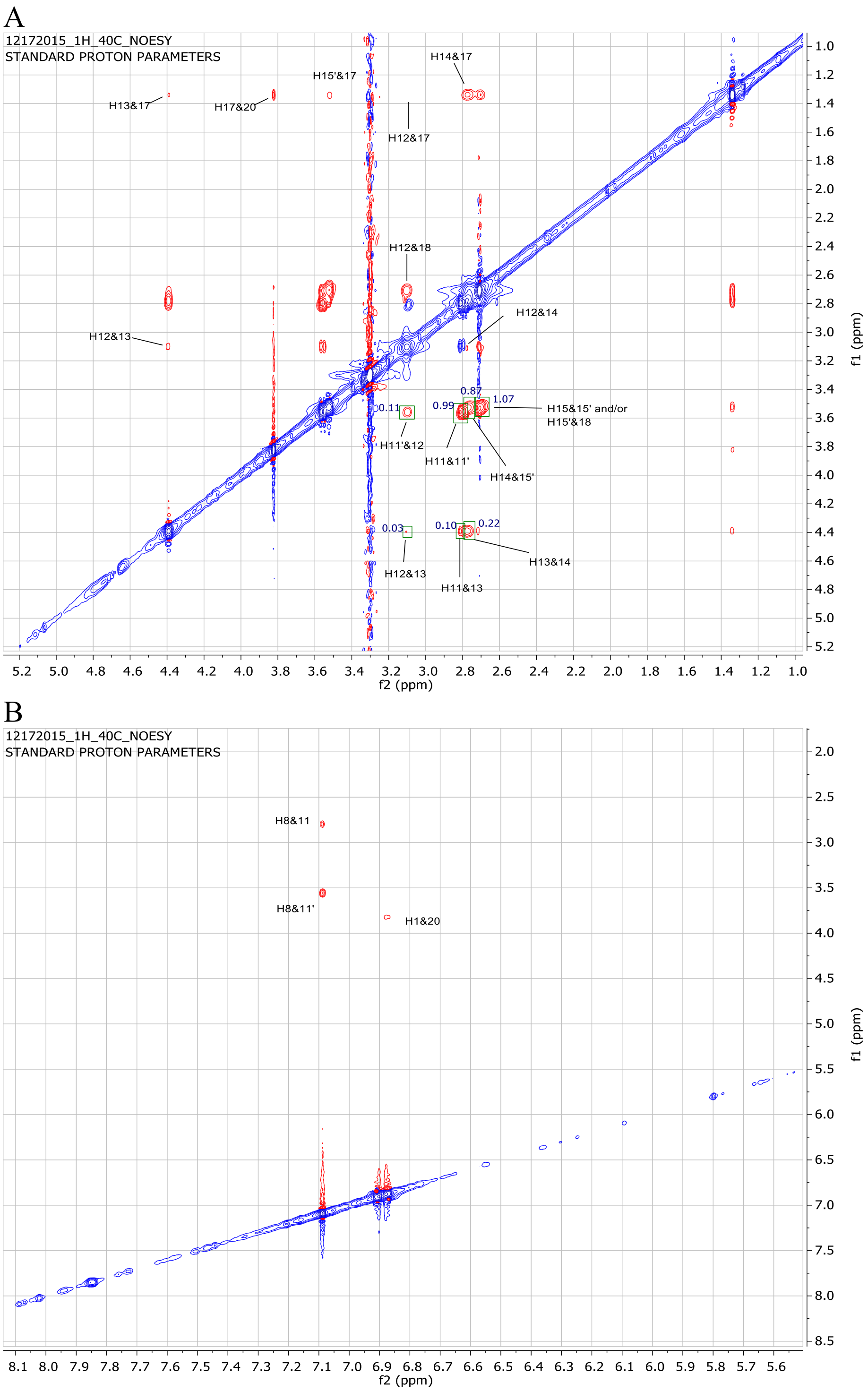


Fig. S7. Nuclear Overhauser effect spectroscopy (NOESY) NMR spectrum of SPF. (*A*) Aliphatic region. The intensities of selected cross-peaks were integrated using Mnova software and shown in the spectrum. (*B*) Aromatic region. Results of the NOESY experiment support the final structures (Fig. 2G and H) due to the presence of NOE signal between H (δH 3.81) and H (δH 1.34), which could only be observed between protons with short spatial distance. For protons on the three consecutive chiral centers, H (δH 4.40) had an intense cross-peak with H (δH 2.77), while a weak signal was observed between H (δH 4.40) and H (δH 3.10) and no signal was observed between H (δH 2.77) and H (δH 3.10). This suggests that H (δH 4.40) and H (δH 2.77) are close to each other and both are distant from H (δH 3.10), which corresponds to (*R, S, S*) or (*S, R, R*) configuration on C 12, 13, 14 (δC 76.7, 88.2 and 37.2). This was further supported by NOESY signals between H (δH 2.79, 3.56) and the three H on chiral centers. H (δH 4.40) had a cross-peak with H (δH 2.79) but no cross-peak with H (δH 3.56). However, the opposite was observed for H (δH 3.10) which had a cross-peak with H (δH 3.56) but no cross-peak with H (δH 2.79).

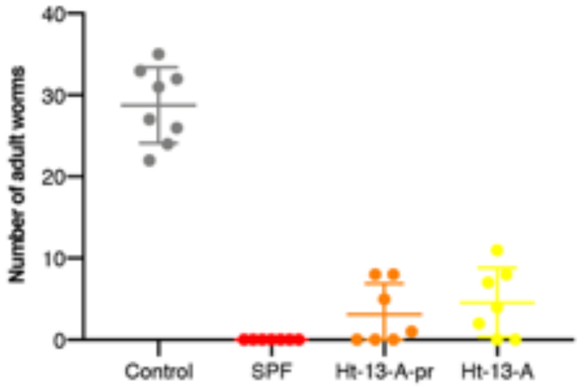


Fig. S8. Stringent mouse infection experiment. Numbers of adult worms recovered from mice exposed to ~100 cercariae that were pre-treated with APW (N=8), 2.5 µM SPF (N=7), 2.5 µM Ht-13-A (N=7) or 2.5 µM Ht-13-A-pr (N=7). The mouse tail was lifted slightly during exposure so that its tip was 1-2 cm from the bottom of the test tube, avoiding direct contact with paralyzed cercariae. Data are mean ± S.D.

Table S1. Summary of protons and carbons from $^1$H, COSY, HSQC, HMBC and NOESY.

| | $^{13}$C | | | $^1$H | | | | | |
|---|---|---|---|---|---|---|---|---|---|
| Position | $\delta_C$ (detected) | $\delta_C$ (predicted) | mult. | $\delta_H$ | Peak area | mult. | COSY | HMBC | NOESY |
| 1 | 116.8 | 110.4 | CH | 6.86 | 1.12 | d (*J*=8.6 Hz) | 6 | 3, 5 | 18 |
| 2 | 143.1 | 148.2 | C | | | | | | |
| 3 | 143.7 | 143.1 | C | | | | | | |
| 4 | 120.6 | 120.3 | C | | | | | | |
| 5 | 138.1 | 132.5 | C | | | | | | |
| 6 | 106.6 | 103.7 | CH | 6.90 | 1.00 | d (*J*=8.6 Hz) | 1 | 2, 4 | |
| 7 | 110.6 | 110.8 | C | | | | | | |
| 8 | 124.9 | 123.0 | CH | 7.09 | 1.00 | s | | 4, 5, 7 | 11,11’ |
| 9 | | | NH | | | | | | |
| 10 | | | O | | | | | | |
| 11 | 29.0 | 32.5 | $CH_2$ | 2.79 | | Overlap | 11’, 12 | 7, 8,12,13 | 8,11’,13 |
| 11’ | 29.0 | 32.5 | $CH_2$ | 3.56 | 1.04 | dd (*J*=14.3 Hz, 3.8Hz) | 11, 12 | 4,7,8,12,13 | 8,11,12 |
| 12 | 76.7 | 72.1 | CH | 3.10 | 1.16 | br | 11, 11’, 13 | | 11’,13,14,17,18 |
| 13 | 88.2 | 84.5 | CH | 4.40 | 1.05 | dd (*J*=9.4 Hz, 6.8Hz) | 12, 14 | 3,11,15 | 11,12,14,17 |
| 14 | 37.2 | 38.9 | CH | 2.77 | | Overlap | 13, 15’, 17 | 15,17 | 12,13,15’,17 |
| 15 | 65.8 | 62.6 | $CH_2$ | 2.70 | | Overlap | 15’ | 12 | 15’,17 |
| 15’ | 65.8 | 62.6 | $CH_2$ | 3.52 | 1.08 | dd (*J*=9.6 Hz, 6.8Hz) | 14 | 12,13 | 14, 15,17,18 |
| 16 | | | N | | | | | | |
| 17 | 14.0 | 14.6 | $CH_3$ | 1.34 | 3.01 | d (*J*=6.8 Hz) | 14 | 13, 14, 15 | 12,13,14,15,15’,20 |
| 18 | 41.7 | 43.7 | $CH_3$ | 2.71 | | s | | 12, 15 | 12,15’ |
| 19 | | | O | | | | | | |
| 20 | 62.2 | 56.1 | $CH_3$ | 3.81 | 2.98 | s | | 2 | 17 |
